# Shared genetic architecture links energy metabolism, behavior and starvation resistance along a power-endurance axis

**DOI:** 10.1101/2024.09.05.611396

**Authors:** Berra Erkosar, Cindy Dupuis, Loriane Savary, Tadeusz J. Kawecki

**Affiliations:** Department of Ecology and Evolution, Faculty of Biology and Medicine, University of Lausanne, Switzerland; FIND, Geneva, Switzerland

**Keywords:** experimental evolution, dietary adaptation, physiology, metabolome, Drosophila, trait syndromes, variational modules, genetic covariance

## Abstract

Shared developmental, physiological and molecular mechanisms can generate strong genetic covariances across suites of traits, constraining genetic variability and evolvability to certain axes in multivariate trait space (“variational modules” or “syndromes”). Such trait suites will not only respond jointly to selection; they will also covary across populations that diverged from one another by genetic drift. We report evidence for such a genetically correlated trait suite in *Drosophila melanogaster*. It links high expression of glycolysis and TCA cycle genes, high abundance of mitochondria and high spontaneous locomotor activity with low degree of adiposity, and low endurance and early death under starvation. This “power-endurance” axis is also aligned with abundance of certain metabolites, notably low trehalose (blood sugar) and high levels of some amino acids and their derivatives, including creatine, a compound known to facilitate energy production in muscles. Our evidence comes from six replicate “Selected” populations adapted to a nutrient-poor larval diet regime during 250 generations of experimental evolution and six “Control” populations evolved in parallel on a standard diet regime. We found that, within each of these experimental evolutionary regime, the above traits strongly covaried along the power-endurance axis across replicate populations which diversified by drift, indicating a shared genetic architecture. The two evolutionary regimes also drove divergence along this axis, with Selected populations on average displaced towards the “power” direction compared to Controls. Aspects of this “power-endurance” axis resemble the “pace of life” syndrome and the “thrifty phenotype”; it may have evolved as part of a coordinated organismal response to nutritional conditions.

## Introduction

Adaptive phenotypically plastic responses to external environmental changes and internal states of an organism often require highly coordinated changes in multiple traits, mediated by changes in expression of many genes. This favors the evolution of regulatory integration of traits through developmental, metabolic and signaling mechanisms that ensure that the responses are coordinated across the traits and genes and expressed at the right time. Once such phenotypic integration of multiple traits has evolved, it may constrain their genetic architecture. On the one hand, it may involve redundance and compensatory mechanisms that would buffer the phenotypes against effects of mutations that affect the expression of single effector genes. On the other hand, genetic variants in key genes regulating such integrated responses would be highly pleiotropic. As a consequence, the traits would become highly genetically correlated, with variation limited to certain axes within the multidimensional phenotypic space, and a substantial fraction of theoretically possible trait value combinations not accessible to evolution, at least in a short term (Wagner and Altenberg, 1996; Gomulkiewicz and Houle, 2009; Blows et al., 2015). Such covarying suits of traits have been found at the level of gene expression (e.g., Blows et al., 2015), life history (e.g., Promislow and Harvey, 1990; Bengston, 2018; Arnqvist et al., 2022) and behavior (e.g., Royauté et al., 2015); they are referred to as variational modules (Wagner and Altenberg, 1996; Blows et al., 2015) or trait syndromes (Dammhahn et al., 2018).

Here we report the inadvertent discovery of a syndrome of traits linking aspects of gene expression, metabolome, abundance of mitochondria, adiposity, behavior and starvation resistance, in *Drosophila melanogaster*. We investigated these traits in a set of six “Selected” populations that, in the course of >250 generations of experimental evolution, became adapted to a nutritionally very poor larval diet, and in six “Control” populations of the same origin that evolved in parallel on a standard larval diet.

Adaptation of the Selected populations is evidenced by their improved larval growth and better survival on the poor diet (Kolss et al., 2009); it is associated with wide-ranging changes in gene expression, nutrient acquisition and larval metabolism (Erkosar et al., 2017; Cavigliasso et al., 2020; Cavigliasso et al., 2023; Cavigliasso et al., 2024), and with differentiation of SNP frequencies at >100 genomic regions (Kawecki et al., 2021). Even though the poor versus standard diet regime has been applied to larvae (adults of all populations being transferred to fresh standard diet), many of the differences in gene expression and metabolome carry over from the larval to adult stage, likely reflecting lack of life stage specificity of variants favored by the larval selection regimes (Erkosar et al., 2023). Yet, freshly emerged adults of Selected populations are less resistant to starvation (Kawecki et al., 2021) and show lower fecundity than Controls even when both are raised on the poor diet, to which Selected but not Control populations have adapted (Erkosar et al., 2023), implying a cost of larval adaptation for adult performance.

In this study we initially set out to study the consequences of adaptation to larval undernutrition for the dynamics of energy stores of adult flies and their potential contribution to differences in starvation resistance. In an apparently paradoxical plastic response, *Drosophila* raised under nutrient limitation eclose with a greater relative triglyceride (fat) content (Borash and Ho, 2001; Baldal et al., 2005; Klepsatel et al., 2020; Min et al., 2021). This has been interpreted as adaptive plasticity anticipating harsh nutritional conditions after metamorphosis (Klepsatel et al., 2020; Min et al., 2021). If so, this plastic response should have been selected against and thus diminished in the Selected populations because they evolved under nutritional hardship limited to the larval stage – adults were transferred to fresh standard food. Therefore, Selected flies should eclose leaner than Control flies when raised on the poor diet. If confirmed, this could explain why freshly emerged virgin Selected flies are less resistant to starvation than Controls, in particular when raised on the poor diet (Kawecki et al., 2021). Furthermore, we predicted that the Control flies raised on the poor diet would eliminate excess fat over the first few days of adult life on fresh standard diet, resulting in convergence in energy stores between Selected and Control flies. If differences in energy stores were the main reason for the difference in starvation resistance between freshly emerged Selected and Control flies, this difference should strongly diminish following convergence in energy stores over the first few days of adulthood. To test these predictions, we quantified triglycerides (and the other main energy store glycogen) in freshly emerged and in 4-day old Selected and Control flies and measured starvation resistance of 4-day old flies.

Our results supported the predictions about triglyceride content, not about starvation resistance, implying that differences in starvation resistance between Selected and Control flies cannot be explained by differences in energy stores. We therefore hypothesized that it may be due to different potential for energy expenditure. To test this hypothesis, we quantified the mitochondrial DNA copy number relative to nuclear genomes and the activity of citrate synthase, a limiting enzyme in the TCA cycle, as well as re-assessing previously published RNAseq data for the expression of genes involved in energy generation (glycolysis and TCA cycle). To look further for clues to explain differences in starvation resistance, we also compared the abundance of 113 core metabolites in Selected and Control flies subject to 24 h of starvation. Finally, as a behavioral indicator of energy expenditure, we compared behavioral activity of flies and their endurance under food deprivation.

Thus, our initial focus of was on differences between Selected and Control populations, and we found that most of the traits we quantified indeed had diverged between them in the predicted directions.

This does not necessarily imply that these traits share genetic architecture; different environments can select for entire suites of traits even if the traits are not genetically correlated. However, we also found that these traits strongly covaried across replicate populations subject to the same experimental selection, which diversified by genetic drift. Furthermore, the principal axis of this covariance in multidimensional trait space was parallel to the divergence driven by the different dietary selection regimes. Such correlated genetic divergence is not expected to arise by drift acting on genetically independent traits. Thus, our results imply that these traits share genetic architecture, with strong covariance along a principal axis that we refer to as a “power-endurance” axis or syndrome.

## Methods

### Experimental flies

The history of the six Selected and six Control populations is described in more detail elsewhere (Kolss et al., 2009; Erkosar et al., 2023). They originated from the same base population adapted to the lab conditions and our standard diet containing 1.25% w/v dry brewer’s yeast, 9% w/v sugars and 5% cornmeal w/v. Over successive generations the Selected populations are raised on a poor larval diet that contains 1/4 of all macronutrients, with Control populations maintained in parallel on the standard diet. Thus, our standard diet already contains less yeast than most typical *Drosophila* lab diets, and the poor diet is extremely nutrient poor, with non-adapted larvae taking more than two weeks to reach pupation. Adults are transferred to fresh standard diet and additionally fed ad libitum live yeast before egg collection for the next generation.

Most experiments were performed on 4-day old (6-day old for locomotor activity) mated and reproducing females, raised on either poor or standard diet and transferred to fresh standard diet within 16 h of eclosion; some measurements were done directly on freshly emerged females. Before all experiments all populations were raised for 2-4 generations on the standard diet to eliminate the effects of (grand)parental environment. For details of diets and fly husbandry, see Supplementary Methods.

### Protein, triglycerides and glycogen content

From each population raised on each diet we collected two samples of five freshly emerged females and, in a separate experimental block, two samples of five 4 days old females. Quantification of protein, triglyceride and glycogen was performed following protocols adapted from Tennessen et al. (2014), for details see Supplementary Methods. The analysis of the two samples for each stage was performed in two batches on two consecutive days. One sample each of 4 days old females from three Selected and three Control populations from either diet was lost due to a technical error.

Triglyceride and glycogen content was normalized to the total protein content of each sample. This assay was performed after 268 generations of experimental evolution.

### Starvation resistance

This assay was performed after 251 generations of experimental evolution on 4 days old females raised on the poor diet. We set up three replicates of 9-11 females each per population; for Control populations C1 and C6 there were enough females only for two replicates each, and for population C1 these only contained 7 and 8 females. Females of each replicate were placed as a group in a vial with agarose (11 g/l) to provide water and monitored several times daily for dead flies (the intervals varied depending on the time of day). The midpoint of interval during which a given fly died was recorded as its time of death. One fly found dead during the first check 6 h after the onset of starvation was excluded from the analysis.

### Metabolome of starving flies

To compare the metabolic state of starving Selected and Control flies, we used a subset of previously published targeted metabolome data obtained (after 230 generations) from carcasses of 4 days old mated females raised identically as in other experiments reported here (Erkosar et al., 2023). (The carcass consists of fat body with the attached abdominal body wall, with parts of embedded Malpighian tubules, tracheae and hemocytes.) We focused here on poor-food raised flies that have been deprived of food for 24 h. The previously reported analysis (Erkosar et al., 2023) focuses on flies in their normal fed conditions and on comparing the difference between fed Selected and Control flies to the response of Control flies to starvation. The data set consists of quantification of 113 core metabolites (normalized to the protein content), with two samples of 10 carcasses (in two blocks) per population. For details of data treatment and scaling see Erkosar et al. (2023).

### Expression of TCA and glycolysis enzyme genes

We used the previously published RNAseq data obtained after 190 generations of selection from carcasses of 4 days old mated females raised on poor and standard larval diet and maintained after eclosion for 3 days on standard diet (Erkosar et al., 2017; Erkosar et al., 2023) (NCBI GEO accession GSE193105). Among the 8701 genes included in that data set we identified 21 genes coding for enzymes catalyzing the reactions of the TCA cycle and 12 genes coding for the enzymes catalyzing the 10 steps of glycolysis. From the original analysis that RNAseq data (Erkosar et al., 2023) we extracted the estimated differences in expression (log_2_ fold-change values) of these genes between the means of Selected and Control populations for flies raised on poor food larval diet. We also used the normalized expression levels of each of these genes for each population (log_2_ counts per million) to use in a PCA.

To test whether the distribution of these differences deviates from what would be expected from an equivalent random set of genes, we first binned the 33 glycolysis and TCA cycle genes by their mean level of expression across all conditions at unit intervals on the log_2_ scale. We next sampled at random 33 genes with the same distribution of mean expression levels among the bins, repeating this 1000 times. We then used the Kolmogorov-Smirnov test to compare the distribution of log_2_ fold change values across the 33 respiratory genes with the distribution of pooled randomized samples.

### Citrate synthase activity and mitochondria quantification

To quantify citrate synthase activity, we collected from each population and each larval diet a single sample of 30 mature female flies (thus the replicate populations are the only level of replication in this assay). We used Citrate Synthase Activity Assay Kit (Sigma, #MAK193) and normalized the reaction rate to the protein content in the sample (for details see Supplementary Methods). The sample from Selected population S4 raised on poor diet was lost. This assay was performed after 245 generations of experimental evolution.

As an attempt to quantify the mitochondria (after 267 generations), we estimated the ratio of mitochondrial to nuclear DNA copy number in samples consisting of thoraxes of five mature females (three replicate samples per population and diet). Samples from two diets were collected on different dates, and the populations within each diet were further split between two days, forming four blocks. DNA was extracted from these samples with the TRIzol reagent (#15596018, ThermoFisher Scientific, Switzerland) following manufacturer’s protocol. We then performed qPCR on two mitochondrial genes, *Cox1* and *ATP6*, and two nuclear genes, *Act42a* and *Rpl32*. The relative amount of mitochondria DNA was calculated as the ratio geometric mean of the concentration of the two mitochondrial genes divided by the geometric mean of the two nuclear genes (for primers and their efficiencies see Supplementary Table S1).

### Locomotor activity

We used the DAM2 Drosophila Activity Monitors by Trikinetics (http://www.trikinetics.com) to quantify locomotor activity of single mature females from Selected and Control populations under conditions of food deprivation. Only flies raised on poor larval diet were used. The system counts the number of times a fly enclosed in a tube crosses an infrared beam. During the test the flies were provided with moisture (agarose) but no food. The flies were introduced in the system within the first hour after lights on (shortly before 9:00); the recording started at 11:00 and continued for 93 h. Because of throughput limitations we split the assay into three block run consecutively, whereby each block included two Selected and Two Control populations. Two activity monitors were used in each block, with individuals from each population split roughly equally between the monitors and positioned randomly within them (12-16 flies in total except for population C5, where only 9 flies were available). This assay was performed after 245 generations of experimental evolution.

Flies in the activity monitors were not monitored for survival. However, with one exception of a fly that last moved 13 h from the onset of recording and was excluded from the analysis, all flies remained capable of movement at least until 36 h from the onset of recording (38 h since they were placed in the apparatus). After that timepoint, an increased number of flies ceased to move and did not resume activity (in terms of beam crossings) until the end of recording, reflecting starvation-induced morbidity and mortality. We therefore decided to focus on the analysis on the first 36 h and used the number of total beam crossings during that interval as a measure of each fly’s cumulative activity. We also recorded for each fly the time at which it moved across the beam for the last time as a readout of its locomotor endurance under starvation.

### Statistical analysis

Statistical analysis was performed with SAS v. 9.4. Glycogen and triglyceride content, citrate synthase activity and the ratio of mitochondrial to nuclear genomes were ratios of two measurements and thus were log-transformed (using log_2_) for the analysis (Houle et al., 2011). The quantity of protein and total activity (the number of beam crossings within 36 h) were also log-transformed as this improved normality of residuals. Starvation resistance (mean time to death per vial) and endurance (timepoint of the last movement in the locomotor activity assay) represented time-to-event type of data. However, they reflect an underlying process of running out of limited resources rather than a kind of Poisson process assumed in survival analysis models, and they did not correspond to the proportional hazard assumption. We therefore analyzed these two traits in with models assuming normal distributions; indeed, their residuals (without data transformation) were reasonably normal.

To test for the differences between populations subject to different evolutionary regimes (i.e., Selected versus Control), these response variables were analyzed with general mixed models (GMM; SAS PROC MIXED) using restricted maximum likelihood (REML). The regime and larval diet on which the experimental flies were raised (poor versus standard) were the fixed factors; for the analysis of protein, glycogen and triglycerides, the stage (i.e. day 1 representing freshly emerged flies versus day 4 representing mature flies) was the third fixed factor. All interactions among the fixed factors were included. Because triglyceride amount showed a complex pattern of 3-way interaction, we additionally fitted separate GMMs to data from day 1 and day 4. Starvation resistance, abundances of metabolites, locomotor activity rate and endurance were only assessed on poor diet-raised flies; hence, regime was the only fixed factor in the analysis. Replicate population (nested in regime) and, where applicable, its interaction with diet and stage, were included as random factors. Experimental batch/block (for locomotor activity and endurance also the activity monitor apparatus, as there were two) was included as a further random factor. Fixed effects were tested with F-tests assuming Satterthwaite approximation for the degrees of freedom (Luke, 2017). Metabolite abundance were adjusted for FDR using the *q*-value approach (Storey and Tibshirani, 2003). Results of the more complex models are reported in supplementary tables; for simpler models, they are included directly in the text.

Activity rate during the first 36 h and the time point of last movement were measured on the same individual flies. This allowed us to explore both individual (within populations) and cross-population relationship between these two traits. To this end, we partitioned the variances and the covariance of the two traits into among- and within-population components (all the time accounting for the difference between regimes as well as the effects of block and apparatus). We estimated the variance components for activity and endurance using SAS PROC VARCOMP, with regime as the fixed factor, and population, block and regime as random factors. We also estimated the same variance components for the sum of the two measurements and obtained the covariance components based on the equation Cov(x,y) = (Var(x+y) – Var(x) – Var(y))/2. The variance and covariance components were then used to estimate among- and within-population correlations.

For metabolite abundance and the expression levels of the TCA and glycolysis genes, we performed a principal component analyses (PCA). For metabolome the PCA was done on estimated population means (Erkosar et al., 2023); for gene expression there was only one sample per population.

Because these variables are measured on a common log scale, we performed the PCA on the covariance rather than correlation matrix (otherwise genes or metabolites with little variability would be given the same weight as those with much variability). We compared the PC scores of Selected and Control populations with a *t*-test.

To explore the correlation structure of within-regime among-population variation, we performed a correlation-based PCA on residuals of population estimates of each trait from the overall means of their corresponding evolutionary regime (i.e., Selected or Control). This analysis included protein content per fly, triglyceride and glycogen content (normalized to protein), mtDNA copy number (all log-transformed), time to death under starvation, activity in 36 h and the timepoint of last movement. We further included the expression of glycolysis and TCA cycle genes and the metabolome of starving flies in the form of residual population scores on their first principal components. (Citrate synthase activity was excluded because of missing estimate for one population.) To estimate to what degree the among-population variation is aligned along the first principal component we used a randomization approach. We performed the PCAs on 100,000 data set in which the values of residuals were permuted at random within each regime independently for each trait. From this, we obtained the “null” distribution of the proportion of variance explained by the first principal component and used it as a refence point to compare the proportion of variance explained by the PCA on real residuals.

To explore which of the metabolic and physiological traits best predict starvation resistance and locomotor activity of poor-diet raised flies, we performed a stepwise selection of GLMs using SAS PROC GLMSELECTION with the selection = stepwise option, using the Bayesian Information Criterion (BIC). This was done on population least-square means of trait values (adjusted for block effects or missing values and back-transformed to the natural scale). The explanatory variables included in this analysis were the evolutionary regime as a categorical variable and the continuous variables: protein content (log-transformed), triglyceride and glycogen content on day 4 (normalized to protein and log transformed), log mitochondrial to nuclear DNA ratio and the first two principal components of metabolome after 24 of starvation. Citrate synthase activity was left out because of missing data for one population.

## Results

### Dynamics of protein and metabolic stores

We compared the amount of soluble protein, glycogen and triglycerides in freshly emerged virgin female flies (collected directly from the diets on which they developed as larvae) and in mature (mated and reproducing) females that had fed on the standard diet for three days. Flies raised on the poor diet contained less than half of the amount of protein of flies raised on standard diet (Fig. 1A; main effect of diet: *P* < 0.0001; see Supplementary Table S2A for details of the statistical analysis). This reflects the stunting effect of the poor diet, already reported in terms of body weight (Kolss et al., 2009). Selected flies contained slightly less protein than Control flies (main effect of evolutionary regime: *P* = 0.028). The protein content did not detectably change between the two time points (*P* = 0.95), suggesting it indeed reflects the structural body size.

**Figure 1.**
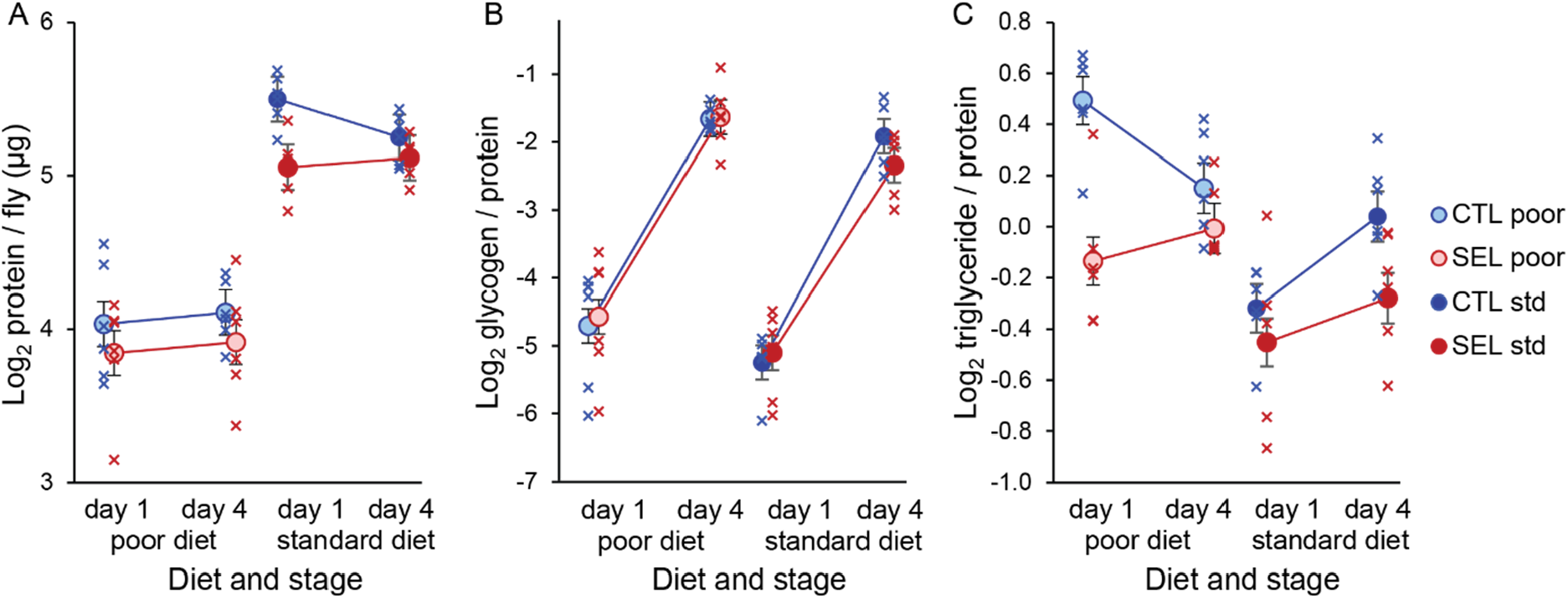
Protein content and energy stores of adult females from Selected and Control populations raised on either larval diet, measured within hours of eclosion (day 1) or after 3 days of mating and feeding on standard diet (day 4). (A) The amount of soluble protein per fly. (B) The quantity of glycogen relative to the protein content. (C) The amount of triglyceride relative to the protein content. Crosses represent population estimates, circles regime means, error bars correspond to standard errors estimated from GMM. *N* = 2 samples of 5 flies per population, diet and stage; several samples were lost (see Methods).

All flies emerged with depleted glycogen stores, which they greatly replenished (more than 10-fold) during the subsequent three days of feeding (Fig. 1A, *P* < 0.0001; Supplementary Table S2B).

Glycogen content did not detectably differ between the Selected and Control populations (*P* = 0.91), but at both timepoints it was about 10% higher in flies raised on poor than standard diet (*P* = 0.0015); no interactions were detected (all *P* > 0.25; Supplementary Table S2B).

The dynamics of triglyceride stores was more complex (Fig. 1C), as indicated by the three-way stage × diet × regime interaction (*P* = 0.0055; Supplementary Table S2C; “stage” refers to freshly emerged versus 4 day old flies). At emergence (i.e., on day 1) Control flies raised on the poor diet had accumulated more triglycerides in relation to their protein content than Control raised on standard diet and Selected flies irrespective of diet (Supplementary Table S3). Selected flies raised on poor diet had more triglycerides than Selected raised on standard diet (although only with *P* = 0.066 after correction for multiple comparisons), but the diet effect was much smaller than for the Control flies (regime × diet interaction on day 1, *P* = 0.0097, Supplementary Table S3).

Over the next three days following the transfer to standard adult diet the triglyceride content of the poor diet-raised Control flies declined (pairwise contrast *F*_1,21.8_ = 6.3, *P* = 0.020) whereas that of Controls on standard diet increased (*F*_1,21.8_ = 7.0, *P* = 0.015). Triglyceride content of Selected flies also tended to increase irrespective of diet (*F*_1,16.4_ = 1.6, *P* = 0.22). As a consequence, 4 days after emergence the triglyceride content was less divergent than on day 1. The remaining difference was mainly due to Selected flies raised on standard diet having somewhat lower triglyceride content than the other three regime × diet combinations (adjusted *P* < 0.05), with the latter being very similar to one another (Supplementary Table S3). In particular, the mean triglyceride content of flies from each regime under conditions to which they were adapted (i.e., Selected raised on poor larval diet, Control raised on standard diet) was nearly identical.

While the spread of the points for individual replicate populations in Figure 1 overestimates the variation among true population means (see Discussion), the general mixed models attributed 50-70% of variance not explained by the fixed factors to the random factors of population and its interactions with stage and/diet (Supplementary Tables S2D, S3C). This indicates substantial genetic diversification among replicate populations even though its phenotypic impact was largely inconsistent between diets and between freshly emerged and mature flies.

### Starvation resistance and starvation metabolome of mature females

We previously found that freshly emerged Selected virgin flies were more susceptible to starvation, dying on average 25% sooner than Controls (Kawecki et al., 2021). Evolution of enhanced starvation resistance in *Drosophila* is often mediated by increased triglyceride stores (Rion and Kawecki, 2007). Thus, those differences in starvation resistance of freshly emerged flies might have been mediated at least in part by the large differences in the amount triglyceride stores between freshly emerged Selected and Control flies. If so, the difference in starvation resistance should vanish following the near-convergence of triglyceride stores during three days of adult feeding. To test this prediction, we compared Selected and Control populations with respect to starvation resistance of mature mated females raised on poor larval diet, after they have fed for 3 days on standard diet. Contrary to our prediction, Selected females died on average 27.5 h (23%) earlier than Control females (Fig. 2A; GMM *F*_1,10.1_ = 17.9, *P* = 0.0017; mean time to death ± SE 93.9 ± 4.6 versus 121.4 ± 4.6 h). We noted that the survival curve of Control population C4 (the leftmost blue line in Figure 1A) actually grouped with Selected populations. This is also the population with the lowest triglyceride content of all Control populations on day 4 (not on day 1), a part of a pattern we return to below.

**Figure 2.**
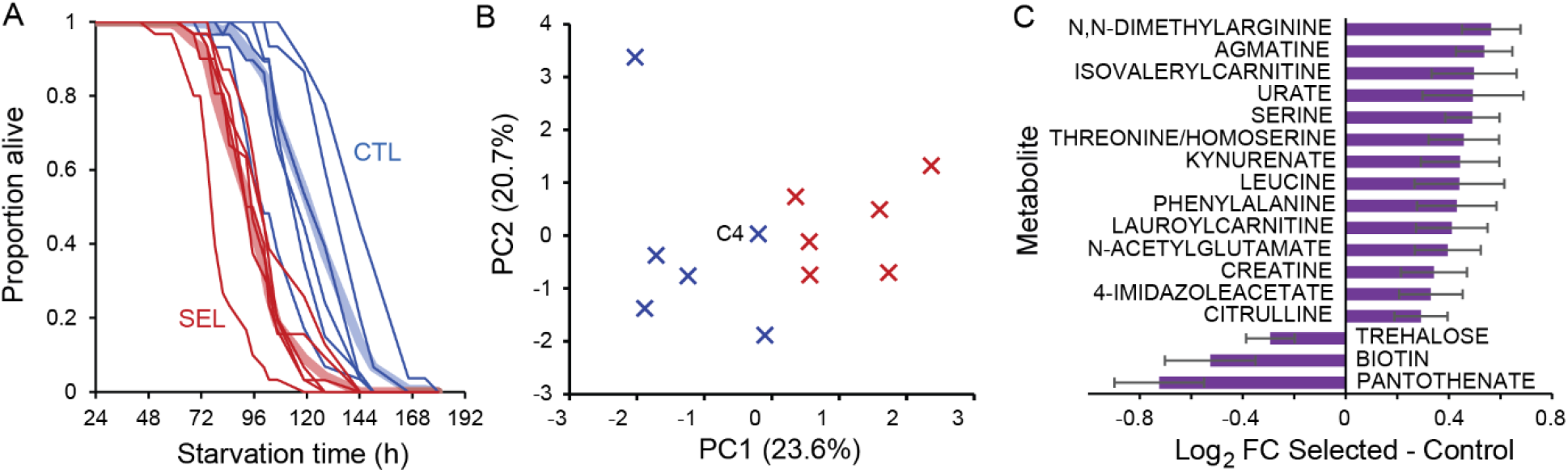
Starvation resistance and starvation metabolome of 4 days old flies raised on poor larval diet. (A) Survival curves of mated females raised on poor diet and maintained on standard diet for 3 days after eclosion and subsequently deprived of food. Fine lines correspond to individual populations, thick paler lines to the means of Selection (red) and Control (blue) populations. *N* = 2-3 replicate vials with 7-12 flies per population. (B) Population scores on the first two principal components based on abundance of 113 core metabolites. (C) Metabolites differentially abundant (*q* < 0.1) between Selected and Control populations in flies subject to 24 h starvation. *N* = 2 samples of 10 fly carcasses per population.

To explore metabolic correlates of these differences in starvation resistance, we compared the abundance of 113 core metabolites in poor-diet raised flies after they have been subject to 24 h of starvation (Supplementary Table S4). Selected and Control populations were clearly differentiated along the first principal component axis based on all 113 compounds (Figure 2B; *t*_10_ = 5.0, *P* = 0.0006), whereby Control population C4 was again close to the Selected populations. Examination of the 17 differentially abundant compounds (Figure 2C) indicates that, compared to starving Control flies, starving Selected flies had lower levels of the main circulating sugar trehalose and a couple of vitamins that are sources of enzymatic cofactors. The compounds with higher abundance in Selected flies include several amino acids and products of amino acid degradation, including urate (uric acid), the main nitrogenous waste product, as well as creatine, an amino acid derivative which (at least in mammals) promotes ATP production in muscles.

### Expression of TCA cycle and glycolysis genes

Earlier death upon starvation could be mediated in part by greater energy expenditure. To compare the potential of Selected and Control flies to generate energy we first examined a previously published genome-wide RNAseq data set from carcasses of 4 days old females (Erkosar et al., 2023) for differences in expression of 33 genes coding for enzymes involved in glycolysis and the TCA (Krebs) cycle. Only one of these 33 genes passed the 5% FDR threshold for expression difference between Selected and Control populations in the original genome-wide expression analysis (Erkosar et al., 2023). However, 30 of the 33 genes had a nominally higher expression estimate in Selected than in Control flies raised on poor larval diet (Figure 3A). For 28 of these genes the estimated expression difference was in a narrow band of log_2_ fold-change of 0.1 to 0.4. The distribution of these expression difference estimates is distinct from a distribution of analogous estimates for a randomly selected subset of genes, the latter sampled in a way to preserve the distribution of overall mean expression levels (Supplementary Figure S1; Kolmogorov-Smirnov test *D*_84_ = 0.68, *P* < 0.0001). The prominent exception – *Scs*β*G* – does not in fact contradict the pattern. It encodes one of two alternative versions of subunit β of succinyl-coA synthetase. This subunit determines whether the reaction will phosphorylate GDP or ADP; a higher *Scs*β*A / Scs*β*G* promotes ATP synthesis at the expense of GTP synthesis (Lambeth et al., 2004).

**Figure 3.**
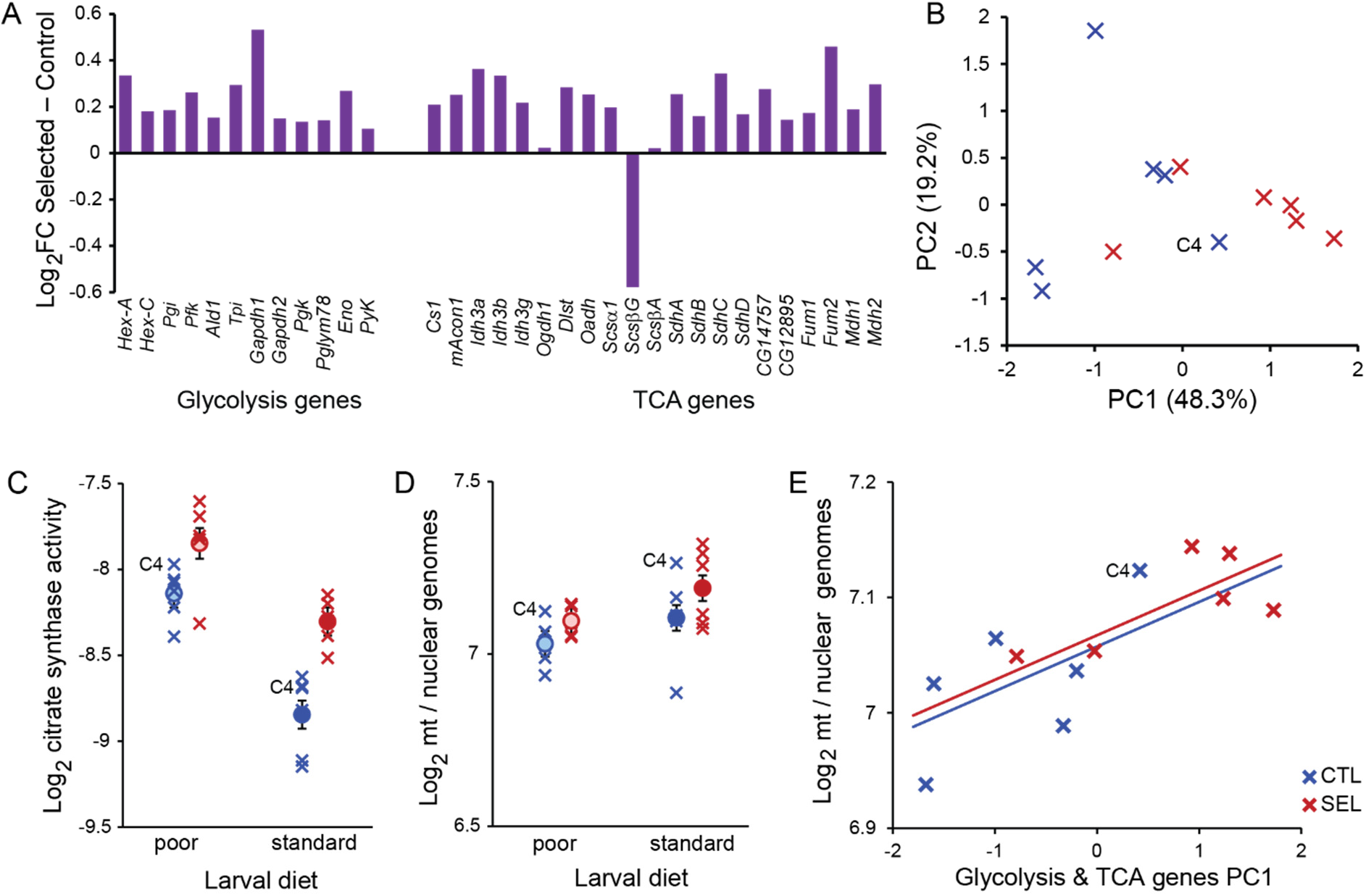
Traits related to energy generation. (A) Estimates of difference between Selected and Control flies in expression of genes for enzymes catalyzing reactions of glycolysis and the TCA cycle, from previously published RNAseq data (Erkosar et al., 2023). (B) PCA on genes from panel A. (C) Activity rate of citrate synthase, the enzyme catalyzing the limiting step of the TCA cycle (in arbitrary units, normalized to protein). *N* = 1 sample of 30 flies per population and diet; the sample for population S4 on poor diet is missing. (D) The ratio of mitochondrial to nuclear genomes in fly thoraxes, based on qPCR on two mitochondrial and two nuclear genes. *N* = 3 replicates of 5 flies per population and diet. (E) The relationship between population scores on PC1 from panel B and the relative quantity of mitochondria. The lines are predictions from ANCOVA in Supplementary Table S7. Circles represent regime means ± SE; crosses correspond to population-specific estimates. Control population C4 is indicated.

Even though PCA on these 33 genes did not perfectly separate the Selected and Control populations, the two regimes differed significantly along the first PC axis, with population C4 again deviating most in the direction of Selected (Figure 3B, *t*_10_ = 2.8, *P* = 0.018). PC1 scores calculated separately for glycolysis and TCA genes were highly correlated across populations (*r* = 0.88, *P* = 0.0002; Supplementary Figure S2), implying these two sets of genes respond in a coordinated way.

Therefore, while the power to detect differences of this magnitude for any particular gene in genome-wide expression data is very weak, pooled evidence from glycolysis and TCA cycle genes suggests that the capacity for sugar catabolism and ATP generation is on average somewhat elevated in Selected flies compared to Controls when both are raised on poor diet.

We previously reported that Selected – Control gene expression differences in adult flies are broadly positively correlated with the corresponding difference in larvae. However, this is not the case for the glycolysis and TCA genes, which in the larvae show a variable pattern but mostly tend to be less expressed in Selected than Control populations (Supplementary Figure S3).

### Citrate synthase activity and mitochondrial DNA copy number

To corroborate the hypothesis about a higher capacity of Selected flies for ATP generation, we quantified the catalytic activity of citrate synthase in whole 4 days old female flies. This assay measures the rate of the reaction under unlimited substrate supply, and thus indirectly quantifies the amount of the enzyme; the actual rate of reaction *in vivo* may be limited by supply of the substrate. Citrate synthase is at the entry point of carbon into TCA cycle, and is the limiting step in the cycle; thus, citrate synthase is sometimes referred to as the “pacemaker” of the TCA cycle (Wiegand and Remington, 1986). We found that citrate synthase activity was on average about 50% higher in flies raised on the poor diet (Figure 4; *P* < 0.0001), and about 30% higher in Selected than Control flies irrespective of diet (*P* < 0.0001), with no detectable interaction (*P* = 0.15, Supplementary Table S5). This is consistent with the Selected flies having a greater capacity to generate ATP.

**Figure 4.**
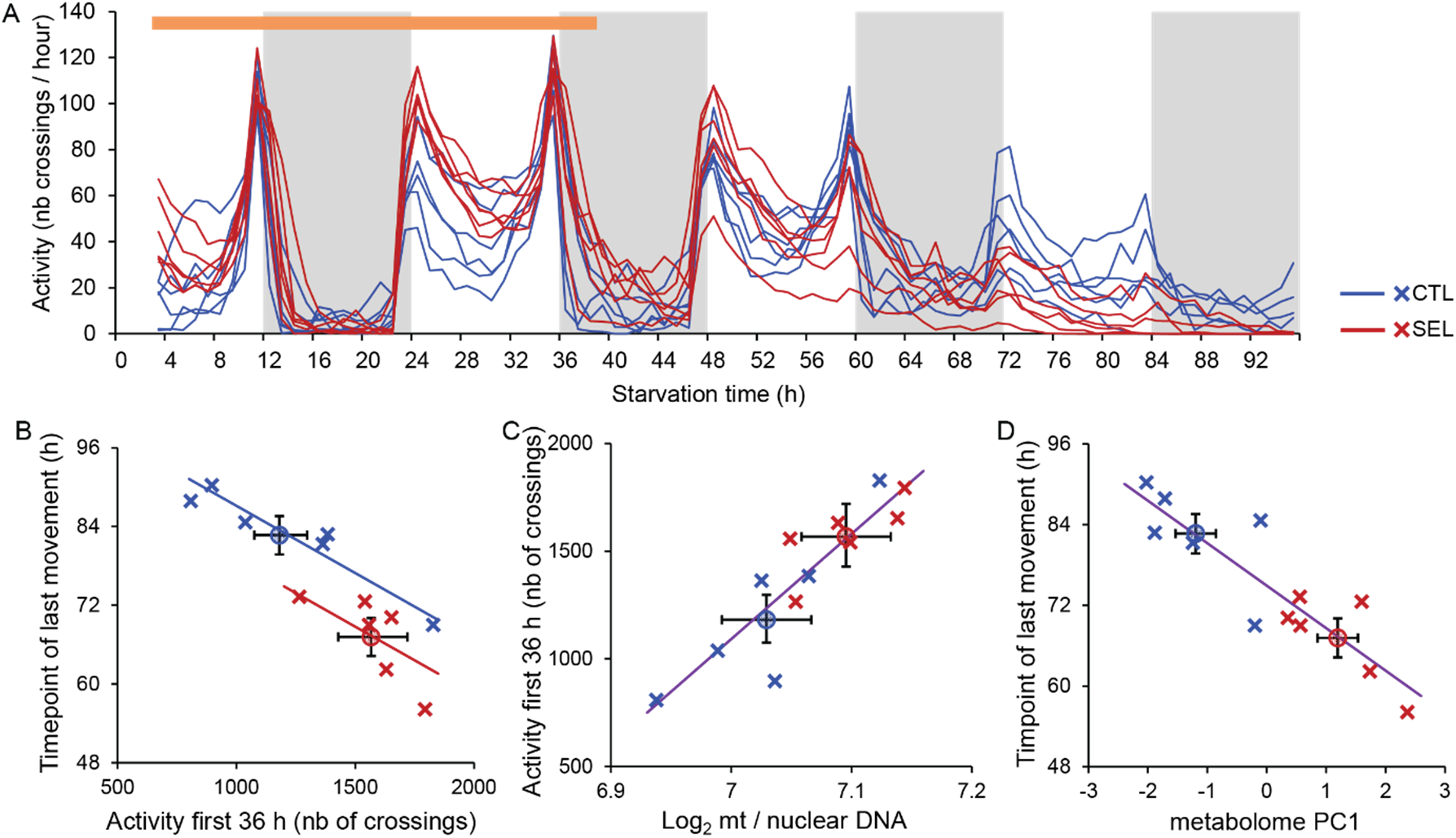
Locomotor activity of starving flies and its relationship with predictive metabolic variables. (A) The pattern of activity (number of beam crossings per hour) over 96 h of food deprivation. The recording began after a 2-h acclimation period. Each line corresponds to a replicate population mean. All flies remained active (i.e., alive) during the first 38 h of experiment, subsequent declines in activity include flies that stopped moving and may have been dead. The orange bar indicates the 36h period used to calculate total activity. (B) Total activity within first 36 h of recording and the timepoint of last recorded movement, and the relationship between population means of the two traits. (C) The relationship between the quantity of mitochondria and total activity. (D) The relationship between the first principal component of the metabolome and the timepoint of last movement. Crosses correspond to means of replicate populations, circles to means of the two regimes; error bars represent SE. The fitted lines in (B) are predictions from ANCOVA, in (C) and (D) they are regression lines fitted jointly to all 12 population means. *N* = 12-16 flies per population, except C5 where *N* = 9.

As a complementary approach to assess if Selected and Control populations differ with respect to energetic potential, we compared the mtDNA genome copy number relative to the nuclear genomes. We did this on thoraxes because this section of the insect body contains mostly muscles, including those involved in walking and flying; muscles are particularly rich in mitochondria and generate large quantities of ATP. In contrast to citrate synthase activity, the ratio of mitochondrial to nuclear genomes was smaller in flies raised on poor diet than on standard diet (*P* = 0.015, Supplementary Table S6). While there was a trend for the mtDNA copy number to be greater in Selected than Control flies, it was not significant (*P* = 0.11), nor was there any hint of interaction (*P* = 0.74). Population C4 again stood out from among Control populations with respect to the relative amount of mitochondria in the direction of the Selected populations’ values (Figure 3D). If this population were removed, the difference between Selected and Control populations would become significant (*F*_1,10.1_ = 6.6, *P* = 0.030). It should be noted that the range of variability for the mitochondrial to nuclear genome ratios was much smaller than for citrate synthase activity (total range of population means on log_2_ scale 0.43 and 1.54, respectively). Nonetheless, within each diet treatment these two measures were moderately positively correlated across populations (poor diet: *r* = 0.64, standard diet: *r* = 0.44). More strikingly, the mtDNA copy number in flies raised on poor diet was quite highly correlated with the first principal component of the glycolysis and TCA cycle gene expression (*r* = 0.77). When included as a covariate, PC1 “explained” virtually all of the difference between regimes (Figure 3E, Supplementary Table S7). It should be noted that the low within-population replication of gene expression, citrate synthase and mitochondria quantity means that the estimates of population values are likely noisy; furthermore, they were taken dozens of generation of selection apart (see Methods). The fact that they are nonetheless correlated not only supports the hypothesis that the Selected flies invest more in the machinery of ATP generation; it also suggests that replicate populations vary in this respect in a consistent way.

### Locomotor activity and endurance upon starvation

The results reported above suggest a higher potential for ATP generation by Selected than Control flies, and locomotion is among the most energetically costly physiological processes. We therefore quantified locomotor (walking) activity of individual mature females under food deprivation, focusing on flies raised on the poor larval diet. Over the first two days fly activity showed the typical pattern of peaks early and late in the day, moderate activity in between and low activity at night. This pattern subsequently degenerated, in part likely because many flies ceased moving, being morbid or dead due to starvation (Figure 4A). For reasons explained in Methods, we based our analysis on total cumulative activity within the first 36 h of recording; except for one individual removed from the analysis all flies remained alive throughout this period. While Selected flies tended to show a higher activity than Controls, this difference was just above the 5% significance threshold (Figure 4B, horizontal axis; *F*_1,8.9_ = 5.0, *P* = 0.052, Supplementary Table S8). The pattern was again broken by Control population C4, which had the highest cumulative activity estimate of all twelve populations (Figure 4B); if this population were removed, the difference between regimes would become significant (*F*_1,8.3_ = 11.6, *P* = 0.0088).

From the activity monitoring data we also derived the timepoint (hour) of the last movement recorded for each fly. This can be interpreted as an estimate of endurance, albeit a highly imperfect one because many flies remained active until the end of the 96-hour recording period. Flies from Selected populations stopped moving on average about 15.5 hours (18%) earlier than those from Control populations (*F*_1,9.8_ = 14.8, *P* = 0.0034, Supplementary Table S8), even though population C4 again grouped with Selected populations (Figure 3B vertical axis). This likely underestimates the true difference in how long the flies can move because only 2 out of 91 Selected flies moved in the last 3 h of the recording, compared to 25 out of 82 Control flies.

We found a strong negative correlation between population means of total activity in the first 36 h and the timepoint of last activity. Based on (co)variance component estimates, the among-populations within-regime correlation was *r* = –0.95, whereas the within-population correlation of these two traits was only *r* = –0.18 (Supplementary Table S9), implying a strong genetic covariance. Consequently, when a model with regime as a fixed factor and mean activity as a covariate was fitted to population means of endurance, the activity-adjusted difference between Selected and Control populations was reduced to 8.2 h but remained significant (fitted lines in Figure 3B; regime *F*_1,9_ = 9.5, *P* = 0.013, mean activity *F*_1,9_ = 24.9, *P* = 0.0007). Thus, a substantial part but not all of the difference in endurance between Selected and Control populations can be attributed to the difference in their rate of locomotion during the first 36 h.

### Relationships among traits across populations

The general mixed models returned positive estimates of variance attributable to replicate populations within the two evolutionary regimes (or population × diet interaction) (Supplementary Tables S2D, S3C, S5B, S6B, S8B). To explore the correlation structure of this within-regime among-population variation, we performed PCA on residuals of population values from their respective regime means.

Using residuals renders this analysis independent of differences between overall means of Selected and Control populations. The first principal component of this analysis (Figure 5A) explained 43% of variation, substantially more than the ∼30% that would have been expected if the residuals had been uncorrelated (*P* = 0.0015 based on randomization; Supplementary Figure S4). Expression of glycolysis genes, mtDNA copy number, metabolome of food-deprived flies and locomotor activity in 36 h loaded positively on the first PC whereas the loadings of triglyceride content, starvation resistance and the timepoint of last movement were negative (Figure 5B). The sign of these loadings corresponds one to one to the sign of estimated differences between Selected and Control regimes. Thus, replicate populations – whether Selected or Control – that deviated from their regime mean towards higher expression of glycolysis and TCA cycle genes also deviated upward with respect to mtDNA content, locomotor activity and PC1 of metabolome assessed after 24 h of food deprivation. For all these traits the estimated mean of Selected populations was above that of Controls (although not significantly so for mtDNA content). The same replicate populations scored lower than average for triglyceride content, starvation resistance and the timepoint of the last move, traits that have lower estimated means in Selected than Control populations. The one trait uncorrelated with that main axis of among-population variation – glycogen content – also showed no hint of difference between Selected and Control flies raised on poor diet. Protein content, which reflects the structural size, loads negatively on PC1, consistent with the trend for Selected flies to be smaller than Controls, but is not aligned with the other vectors (Supplementary Figure 5B). This analysis implies that a large fraction of variation among replicate populations from the same evolutionary regime falls along a single axis, and that this axis aligns with the differences between the overall means of Selected and Control populations.

**Figure 5.**
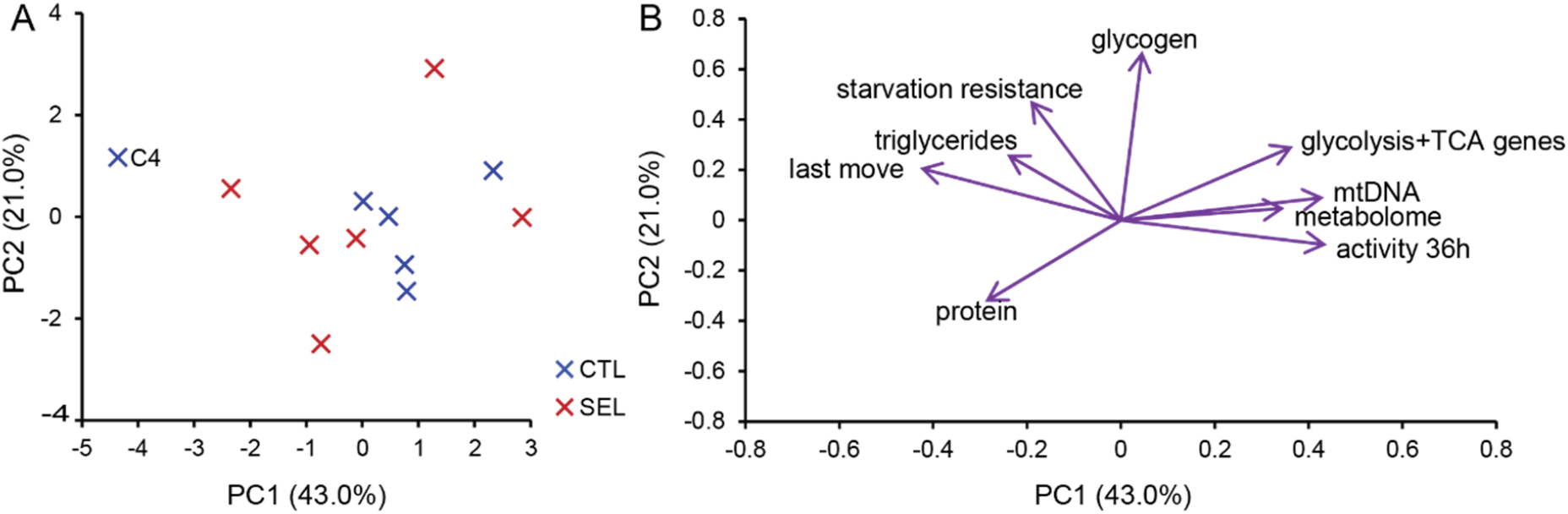
PCA on residuals of population estimates from the means of Selected and Control regimes, for seven phenotypic traits (protein content, triglyceride/protein, glycogen/protein, mtDNA/nuclear DNA copy number, time to death upon starvation, locomotor activity rate and time of last movement) and the first principal components of metabolome under starvation and of glycolysis + TCA cycle gene expression. (A) Population scores in the PCA space. (B) Eigenvectors of the nine traits in the PC1-PC2 space.

We used a model selection approach to explore links between the above metabolic measurements and the variation in starvation resistance, activity and endurance among the populations. The mitochondrial to nuclear genome ratio and the second principal component of metabolome (PC2) were the only variables included in the most parsimonious model for total activity within the first 36 h, “explaining” 91% of variance (Supplementary Table S10). Mt/nuclear genome ratio alone accounted for 79% of variation among populations (Figure 4C). The most parsimonious model for the timepoint of last movement “explained” 88% of variation with metabolome PC1 and mtDNA copy number (Supplementary Table S11), with the PC1 accounting for 79% of variation (Figure 4D). Both those models could nearly perfectly predict the regime means. In contrast, the most parsimonious model for mean time to death under starvation retained regime as the factor accounting for the largest fraction (64%) of variation among population means, with triglyceride content accounting for a further 10% but not passing the significance threshold (*P* = 0.11; Supplementary Figure S5, Supplementary Table S12).

## Discussion

### A plastic response of fat storage to larval undernutrition lost in Selected populations

We set out to investigate the consequences of experimentally evolved adaptation to larval undernutrition for adult metabolic energy stores and their potential contribution to differences in starvation resistance between Selected and Control populations. As expected, we found that, when raised on poor diet, Control flies emerged with >50% higher relative triglyceride content than Selected flies and both Selected and Control flies raised on standard diet. These elevated triglyceride reserves of poor diet-raised Control flies underwent a sharp reduction in the first three days of adult life, nearly to the level observed in Control flies raised on standard diet. Such a counterintuitive phenotypic response – becoming fatter when growing under nutrient (especially protein) shortage – is well known in Drosophila (Borash and Ho, 2001; Baldal et al., 2005; Klepsatel et al., 2020; Min et al., 2021). It has been interpreted as a predictive adaptive response, a form of phenotypic plasticity that primes the individual for poor nutritional conditions in adulthood (Klepsatel et al., 2020; Min et al., 2021).

Consistent with this idea, this response was eliminated in our Selected populations, which evolved under a mismatch between the poor larval and good adult environment. However, such apparently excessive fat accumulation could also result from the larvae having to consume more carbohydrates to accumulate sufficient protein to complete development (Simpson et al., 1993; Gray et al., 2021). Even though the poor and standard diets contained the same protein : carbohydrate ratio, the Selected larvae apparently shifted their digestive physiology towards a higher protein : carbohydrate ratio in absorbed nutrients (Cavigliasso et al., 2020). Whether or not it is adaptive for adults, our results imply that the enhanced triglyceride accumulation is not adaptive in terms of facilitating larval development on poor diet.

Irrespective of the reasons for differences in larval fat accumulation, by day 4 the Selected flies raised on poor larval diet converged to a nearly identical triglyceride content as Control flies raised on standard diet. This result is analogous to evolutionary convergence of triglyceride content of *Drosophila* populations evolving on larval diets with divergent protein : carbohydrate ratio (Gray et al., 2021). It suggests selection towards an optimal level of triglyceride storage in reproductive females, with an optimum that his independent of the larval environment, but also not affected by the 2-fold difference in body size (as measures by protein content) between flies raised on poor versus standard diet.

### Starvation resistance: energy stores versus energy expenditure

Given the convergence of triglyceride content and no difference in glycogen level, we expected that, in contrast to freshly emerged flies, 4 days old Selected and Control flies would have converged in their resistance to starvation. This was not the case. Even though 4 day old flies survived about 50% longer without food than freshly emerged flies (Kawecki et al., 2021), presumably reflecting the replenished glycogen stores, the difference in time to death between Selected and Control flies remained similar: it was 20 h (25%) for freshly eclosed female flies (Kawecki et al., 2021) and 27 h (23%) for 4-day old flies (this study). This implies that energy stores are not a major cause of differences in starvation resistance between Selected and Control flies, at least after they have fed on standard diet for several days.

Rather, multiple lines of evidence point to a potential role of differences in energy expenditure. First, pooled evidence from genes involved in glycolysis and TCA cycle differentiated the Selected and Control populations, with Selected flies showing consistently higher expression estimates of nearly all of these genes than Controls. Second, Selected populations showed a higher citrate synthase activity and tended to have a higher ratio of mitochondrial to nuclear genomes, both indicative of a higher aerobic metabolic capacity (Wiegand and Remington, 1986; Merkey et al., 2011). Third, Selected flies showed higher levels of locomotor activity under food deprivation (although with *P* = 0.052 just above the conventional significance threshold). Taken together, these diverse results suggest that Selected populations have a higher capacity to generate ATP and, based on their higher expression of glycolysis genes, to catabolize sugars. Before we further discuss the implication of these findings, we address the variability among replicate populations within each of the two selection regimes.

### Variation among replicate populations aligns with differences between regimes

Because our initial aim was to understand consequences of adaptation to larval diets, we focused on testing for divergence between Selected and Control populations, in particular when raised on the poor diet, with the six populations in each group being the principal units of replication (Kawecki et al., 2012). However, we found substantial variation among the replicate populations within each evolutionary regime, which rendered the differentiation between the regimes not quite consistent. It should be noted that the expected variance among observed population means of a given trait is the sum of two variance components: variance among true population means and residual variance (due to individual variation, vial effects etc.) divided by the number of within-population replicates. Thus, the low number of replicates within populations (*N* = 2-3 samples for most traits, *N* = 1 for gene expression and citrate synthase) means that the spread of population values around the regime means exceeds the actual range of true population values, likely by a substantial margin.

Nonetheless, for most traits the GMM attributed substantial among variation among replicate populations indicating real genetically-based variation among them.

Given the small *Ne* around 100-120 (Kawecki et al., 2021), it is not surprising that, after 250 generations of independent evolution, replicate populations subject to the same conditions diverged genetically from one another by drift and drift-contingent selection. However, an unexpected finding is that a large fraction of variation among replicate populations from the same evolutionary regime, at least for flies raised on poor diet, fell along a single multi-trait axis (corresponding to the first principal component). Furthermore, this axis aligned with the axis that differentiates Selected from Control populations. We refer to this axis as a “power-endurance” trait syndrome. The “power” end of the spectrum is characterized by high expression of energy metabolism genes, greater quantity of mitochondria and high rate of locomotor activity, but low triglyceride stores, low resistance to starvation and low endurance under starvation. The inverse combination of traits characterizes the “endurance” end. This axis is also aligned with the first PC of metabolome of starving flies, with the “power” end characterized by low trehalose, biotin and pantothenate concentration combined with high levels of some free amino acids and their derivatives. The 43% of multivariate normalized variance explained by this axis probably underestimates the strength of the underlying genetic correlations – the PCA it is based on estimated populations means, which are subject to a substantial measurement error (see above), obscuring genetic correlations between traits. This is exemplified by the relationship between activity rate and endurance, which were measured on the same individual flies, thus permitting partitioning of phenotypic (co)variance, resulting in an estimate of among-populations correlation as high as –0.95.

Differentiation with respect to multiple functionally linked traits between populations subject to different selection pressures is expected even if the traits are genetically independent. However, a correlated differentiation of genetically independent traits is not expected to arise by drift in populations evolving in the same environment, as genetic variants affecting the traits would drift independently. This implies that the power-endurance axis of variation observed both between regimes and among populations within regimes represents an integrated suite of traits sharing genetic architecture and constrained to evolve in concert by drift as well as by selection.

### Power-endurance axis, “pace of life” syndrome and thrifty phenotype

While regions of suppressed recombination (supergenes) can link traits that are developmentally and physiologically independent, a shared genetic architecture is more likely to reflect developmental, physiological and/or molecular links among traits. Such functional links between some of the traits that define the “power-endurance” axis in our results are also supported by other studies. The rate of energy dissipation (i.e., metabolic rate), is positively genetically correlated with spontaneous locomotor activity rate in *Drosophila* (Videlier et al., 2021) and mice (Gębczyński and Konarzewski, 2009). Spontaneous locomotor activity is turn negatively genetically correlated with starvation resistance in *Drosophila* (Harshman et al., 1999; Schwasinger-Schmidt et al., 2012; Everman et al., 2019), as is metabolic rate (Harshman et al., 1999). Also in line with the power-endurance axis, enhanced sexual selection has led to the evolution of higher metabolic rate, higher locomotor activity and lower starvation resistance (albeit no differences in lipid content) in *D. pseudoobscura* males (Garlovsky et al., 2022). At the level of energy metabolism, downregulation of glycolysis genes (Sorensen et al., 2007) and an elevated trehalose level (Schwasinger-Schmidt et al., 2012) were reported as correlated responses to selection for starvation resistance in *D. melanogaster*.

Metabolites more abundant in Selected populations and aligned with the “power” direction of the axis include creatine and citrulline. Both enhance short-term high-intensity exercise capacity in human athletes (Kerksick et al., 2018), and creatine supplement stimulates locomotor activity in *Drosophila* (Kim et al., 2020). Biotin in turn is less abundant in Selected populations and aligns with the “endurance” direction of the axis; consistent with this, *Drosophila* on biotin-deficient diet show higher locomotor activity and lower starvation resistance (Smith et al., 2007). These observations about specific metabolites are anecdotic but tantalizing; while little can be concluded from them about mechanisms that integrate these traits along the power-endurance axis, they further support the existence of such an integrated trait suite.

Aspects of this trait suite resemble aspects of the “pace-of-life syndrome”, which emphasizes links between high metabolic rate and “fast” life history: fast development, high fecundity and short lifespan (Dammhahn et al., 2018; Arnqvist et al., 2022). The Selected populations indeed evolved faster larval development than Controls, mediated in part by faster larval growth on the poor diet and in part by a smaller critical size for metamorphosis initiation (Kolss et al., 2009; Vijendravarma et al., 2012; Cavigliasso et al., 2023). However, they have a lower fecundity, even when raised on poor larval diet (Erkosar et al., 2023). We have argued that much of the divergence between Selected and Control populations in adult gene expression and metabolism is non-adaptive consequence of pleiotropic effects of allele favored by the strong selection imposed by the poor diet on larval physiology (Erkosar et al., 2023). Yet, while there is a general pattern of a positive correlation between differences in larval and adult gene expression, this is not the case for glycolysis and TCA cycle genes. Furthermore, Control population C4, which groups with Selected rather than Control populations on the power-endurance axis for the adult traits, does not deviate from other Control populations for egg-to-adult developmental time, larval gene expression and larval metabolome (Cavigliasso et al., 2023; Cavigliasso et al., 2024). This weakens the hypothesis that the evolution of Selected populations towards high power / low endurance at the adult stage is a byproduct of selection for fast larval development.

There are also some parallels between the “endurance” direction of the trait suite and the “thrifty phenotype” invoked to explain human (and rodent) susceptibility to metabolic diseases such as diabetes: higher blood sugar level, lower metabolic rate, lower propensity to exercise and greater adiposity (Bouchard, 2007; Wells, 2011). Such a thrifty phenotype in mammals is induced by intra-uterine undernutrition and is thought to promote post-natal or adult tolerance to famine that such undernourished pregnancy may foretell. This potentially adaptive plastic response becomes a health problem when the organism encounters plenty of food (Bouchard, 2007; Wells, 2011). Extending this argument, one might speculate that our Control flies raised on the poor diet exhibit an ancestral “thrifty phenotype” response, and that this response had been selected against under the Selected regime, which involves a switch from larval undernutrition to abundant nutrition in the adult stage.

Irrespective of the specific selection pressures that drove the divergence between Selected and Control populations along the “power-endurance” axis, the fact that drift moved populations along the same axis implies a relatively simple genetic architecture linking metabolism, behavior and life history. Presumably this architecture affects key regulatory mechanisms that may have first evolved to generate a coordinated adaptive plastic response of multiple traits to nutritional conditions. Once such an integrated suite of traits, or “syndrome” exists, genetic variation will tend to generate covariance mainly along the principal axis of this syndrome. This will constrain evolutionary change, at least in the short term, in particular if the main axis of covariance does not align with the fitness landscape (Blows et al., 2004).

## Supporting information

Original Data Files

## Data availability

The (previously published) RNAseq data are available at NCBI Gene Expression Omnibus, accession <u>GSE193105</u> (Kawecki et al., 2022).The previously published metabolome data are available as <u>supplementary file 8</u> in Erkosar et al. (2023). The new phenotypic data are available as supplementary material.

## Author contributions

B.E. and T.J.K. designed the research, B.E., C.D. and L.S. performed all experiments, T.J.K. analyzed the data and wrote the article.

## Acknowledgements

We thank Hector Gallart-Ayala, Julijana Ivanisevic and the Metabolomics Unit of the Faculty of Biology and Medicine, University of Lausanne for carrying out the metabolome quantification and advice on metabolome analysis, the Genomic Technologies Facility of the University of Lausanne for the RNAseq data, and Samuel Charberet for comments on an earlier version of the manuscript.

## References

Arnqvist G, Rönn J, Watson C, Goenaga J, Immonen E, 2022. Concerted evolution of metabolic rate, economics of mating, ecology, and pace of life across seed beetles. Proc Natl Acad Sci USA 119:e2205564119. doi: 10.1073/pnas.2205564119.

Baldal EA, van der Linde K, van Alphen JJM, Brakefield PM, Zwaan BJ, 2005. The effects of larval density on adult life-history traits in three species of Drosophila. Mechanisms of Ageing and Development 126:407–416. doi: 10.1016/j.mad.2004.09.035

Bengston SE, 2018. Life-history and behavioral trait covariation across 3 years in Temnothorax ants. Behav Ecol 29:1494–1501. doi: 10.1093/beheco/ary101.

Blows MW, Allen SL, Collet JM, Chenoweth SF, McGuigan K, 2015. The Phenome-Wide Distribution of Genetic Variance. Am Nat 186:15–30. doi: 10.1086/681645.

Blows MW, Chenoweth SF, Hine E, 2004. Orientation of the genetic variance-covariance matrix and the fitness surface for multiple male sexually selected traits. Am Nat 163:329–340. doi: 10.1086/381941.

Borash DJ, Ho GT, 2001. Patterns of selection: stress resistance and energy storage in density-dependent populations of Drosophila melanogaster. J Insect Physiol 47:1349–1356. doi: 10.1016/s0022-1910(01)00108-1

Bouchard C, 2007. The biological predisposition to obesity: beyond the thrifty genotype scenario. International Journal of Obesity 31:1337–1339. doi: 10.1038/sj.ijo.0803610.

Cavigliasso F, Dupuis C, Savary L, Spangenberg JE, Kawecki TJ, 2020. Experimental evolution of post-ingestive nutritional compensation in response to a nutrient-poor diet. Proc R Soc B 287:20202684. doi: 10.1098/rspb.2020.2684.

Cavigliasso F, Savary L, Spangenberg JE, Gallart-Ayala H, Ivanisevic J, Kawecki TJ, 2023. Experimental evolution of metabolism under nutrient restriction: enhanced amino acid catabolism and a key role of branched-chain amino acids. Evolution Letters 7:273–284. doi: 10.1093/evlett/qrad018.

Cavigliasso F, Savitsky M, Koval A, Erkosar B, Savary L, Gallart-Ayala H, Ivanisevic J, Katanaev VL, Kawecki TJ, 2024. Cis-regulatory polymorphism at fiz ecdysone oxidase contributes to polygenic evolutionary response to malnutrition in Drosophila. PLOS Genetics 20:e1011204. doi: 10.1371/journal.pgen.1011204.

Dammhahn M, Dingemanse NJ, Niemelä PT, Réale D, 2018. Pace-of-life syndromes: a framework for the adaptive integration of behaviour, physiology and life history. Behav Ecol Sociobiol 72:62. doi: 10.1007/s00265-018-2473-y.

Erkosar B, Dupuis C, Cavigliasso F, Savary L, Kremmer L, Gallart-Ayala H, Ivanisevic J, Kawecki TJ, 2023. Evolution under juvenile malnutrition impacts adult metabolism and impairs adult fitness in Drosophila. ELife 12:e92465. doi: 10.7554/eLife.92465.

Erkosar B, Kolly S, van der Meer JR, Kawecki TJ, 2017. Adaptation to chronic nutritional stress leads to reduced dependence on microbiota in Drosophila. mBio 8:e01496–01417. doi: 10.1128/mBio.01496-17.

Everman ER, McNeil CL, Hackett JL, Bain CL, Macdonald SJ, 2019. Dissection of Complex, Fitness-Related Traits in Multiple Drosophila Mapping Populations Offers Insight into the Genetic Control of Stress Resistance. Genetics 211:1449–1467. doi: 10.1534/genetics.119.301930.

Garlovsky MD, Holman L, Brooks AL, Novicic ZK, Snook RR, 2022. Experimental sexual selection affects the evolution of physiological and life-history traits. J Evol Biol 35:742–751. doi: 10.1111/jeb.14003.

Gębczyński AK, Konarzewski M, 2009. Locomotor activity of mice divergently selected for basal metabolic rate: a test of hypotheses on the evolution of endothermy. J Evol Biol 22:1212–1220. doi: 10.1111/j.1420-9101.2009.01734.x.

Gomulkiewicz R, Houle D, 2009. Demographic and Genetic Constraints on Evolution. Am Nat 174:E218–E229. doi: 10.1086/645086.

Gray LJ, Sokolowski MB, Simpson SJ, 2021. Drosophila as a useful model for understanding the evolutionary physiology of obesity resistance and metabolic thrift. Fly 15:47–59. doi: 10.1080/19336934.2021.1896960.

Harshman LG, Hoffmann AA, Clark AG, 1999. Selection for starvation resistance in Drosophila melanogaster: physiological correlates, enzyme activities and multiple stress responses. J Evol Biol 12:370–379. doi: 10.1046/j.1420-9101.1999.00024.x.

Houle D, Pelabon C, Wagner GP, Hansen TF, 2011. Measurement and meaning in biology. Quart Rev Biol 86:3–34. doi: 10.1086/658408.

Kawecki TJ, Erkosar B, Dupuis C, Hollis B, Stillwell RC, Kapun M, 2021. The genomic architecture of adaptation to larval malnutrition points to a trade-off with adult starvation resistance in Drosophila. Molecular Biology and Evolution 38:2732–2749. doi: 10.1093/molbev/msab061.

Kawecki TJ, Erkosar B, Dupuis C, Savary L, 2022. Evolutionary and phenotypically plastic response of adult gene expression to larval undernutrition in Drosophila melanogaster. https://www.ncbi.nlm.nih.gov/geo/query/acc.cgi?acc=GSE193105.

Kawecki TJ, Lenski RE, Ebert D, Hollis B, Olivieri I, Whitlock MC, 2012. Experimental evolution. Trends Ecol Evol 27:547–560. doi: 10.1016/j.tree.2012.06.001.

Kerksick CM, Wilborn CD, Roberts MD, Smith-Ryan A, Kleiner SM, Jäger R, Collins R, Cooke M, Davis JN, Galvan E, Greenwood M, Lowery LM, Wildman R, Antonio J, Kreider RB, 2018. ISSN exercise & sports nutrition review update: research & recommendations. Journal of the International Society of Sports Nutrition 15:38. doi: 10.1186/s12970-018-0242-y.

Kim S, Hong K-B, Kim S, Suh HJ, Jo K, 2020. Creatine and taurine mixtures alleviate depressive-like behaviour in Drosophila melanogaster and mice via regulating Akt and ERK/BDNF pathways. Scientific Reports 10:11370. doi: 10.1038/s41598-020-68424-1.

Klepsatel P, Knoblochova D, Girish TN, Dircksen H, Galikova M, 2020. The influence of developmental diet on reproduction and metabolism inDrosophila. Bmc Evolutionary Biology 20. doi: 10.1186/s12862-020-01663-y.

Kolss M, Vijendravarma RK, Schwaller G, Kawecki TJ, 2009. Life history consequences of adaptation to larval nutritional stress in Drosophila. Evolution 63:2389–2401. doi: 10.1111/j.1558-5646.2009.00718.x.

Lambeth DO, Tews KN, Adkins S, Frohlich D, Milavetz BI, 2004. Expression of Two Succinyl-CoA Synthetases with Different Nucleotide Specificities in Mammalian Tissues *. J Biol Chem 279:36621–36624. doi: 10.1074/jbc.M406884200.

Luke SG, 2017. Evaluating significance in linear mixed-effects models in R. Behavior Research Methods 49:1494–1502. doi: 10.3758/s13428-016-0809-y.

Merkey AB, Wong CK, Hoshizaki DK, Gibbs AG, 2011. Energetics of metamorphosis in Drosophila melanogaster. J Insect Physiol 57:1437–1445. doi: 10.1016/j.jinsphys.2011.07.013.

Min KW, Jang T, Lee KP, 2021. Thermal and nutritional environments during development exert different effects on adult reproductive success in Drosophila melanogaster. Ecol Evol 11:443–457. doi: 10.1002/ece3.7064.

Promislow DEL, Harvey PH, 1990. Living fast and dying young: A comparative analysis of life-history variation among mammals. J Zool 220:417–437. doi: 10.1111/j.1469-7998.1990.tb04316.x.

Rion S, Kawecki TJ, 2007. Evolutionary biology of starvation resistance: what we have learned from Drosophila. J Evol Biol 20:1655–1664.

Royauté R, Greenlee K, Baldwin M, Dochtermann NA, 2015. Behaviour, metabolism and size: phenotypic modularity or integration in Acheta domesticus? Anim Behav 110:163–169. doi: 10.1016/j.anbehav.2015.09.027.

Schwasinger-Schmidt TE, Kachman SD, Harshman LG, 2012. Evolution of starvation resistance in Drosophila melanogaster: measurement of direct and correlated responses to artificial selection. J Evol Biol 25:378–387. doi: 10.1111/j.1420-9101.2011.02428.x.

Simpson SJ, Raubenheimer D, Bone Q, 1993. A multi-level analysis of feeding behaviour: the geometry of nutritional decisions. Philosophical Transactions of the Royal Society of London Series B: Biological Sciences 342:381–402. doi: 10.1098/rstb.1993.0166.

Smith EM, Hoi JT, Eissenberg JC, Shoemaker JD, Neckameyer WS, Ilvarsonn AM, Harshman LG, Schlegel VL, Zempleni J, 2007. Feeding Drosophila a Biotin-Deficient Diet for Multiple Generations Increases Stress Resistance and Lifespan and Alters Gene Expression and Histone Biotinylation Patterns123. The Journal of Nutrition 137:2006–2012. doi: 10.1093/jn/137.9.2006.

Sorensen JG, Nielsen MM, Loeschcke V, 2007. Gene expression profile analysis of Drosophila melanogaster selected for resistance to environmental stressors. J Evol Biol 20:1624–1636. doi: 10.1111/j.1420-9101.2007.01326.x.

Storey JD, Tibshirani R, 2003. Statistical significance for genomewide studies. Proc Natl Acad Sci USA 100:9440–9445. doi: 10.1073/pnas.1530509100.

Tennessen JM, Barry WE, Cox J, Thummel CS, 2014. Methods for studying metabolism in Drosophila. Methods 68:105–115. doi: 10.1016/j.ymeth.2014.02.034.

Videlier M, Rundle HD, Careau V, 2021. Sex-specific genetic (co)variances of standard metabolic rate, body mass and locomotor activity in Drosophila melanogaster. J Evol Biol 34:1279–1289. doi: 10.1111/jeb.13887.

Vijendravarma RK, Narasimha S, Kawecki TJ, 2012. Chronic malnutrition favours smaller critical size for metamorphosis initiation in Drosophila melanogaster. J Evol Biol 25:288–292. doi: 10.1111/j.1420-9101.2011.02419.x.

Wagner GP, Altenberg L, 1996. Perspective: complex adaptations and the evolution of evolvability. Evolution 50:967–976.

Wells JCK, 2011. The thrifty phenotype: An adaptation in growth or metabolism? American Journal of Human Biology 23:65–75. doi: 10.1002/ajhb.21100.

Wiegand G, Remington SJ, 1986. CITRATE SYNTHASE: Structure, Control, and Mechanism. Annual Review of Biophysics 15:97–117. doi: 10.1146/annurev.bb.15.060186.000525.

