## Supplementary Figures Tables and Methods for "Shared genetic architecture links energy metabolism, behavior and starvation resistance along a power-endurance axis"

### Supplementary material

Berra Erkosar, Cindy Dupuis, Loriane Savary, Tadeusz J. Kawecki

Department of Ecology and Evolution, Faculty of Biology and Medicine, University of Lausanne, Switzerland

Supplementary Figures S1-S5

Supplementary Tables S1-S9

Supplementary Methods

### Supplementary Figures

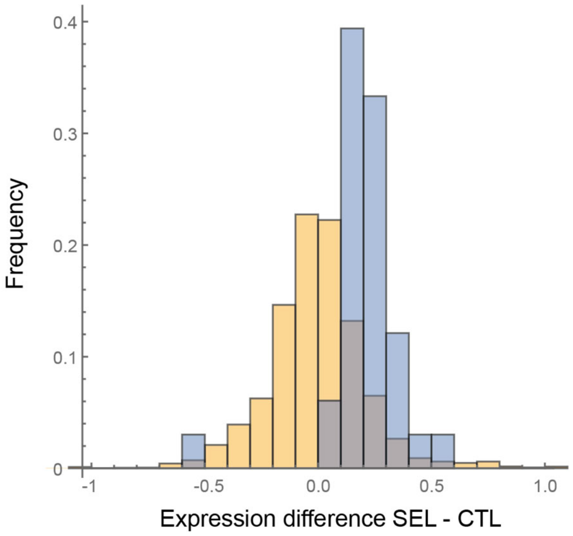


**Supplementary Figure S1**. Glycolysis and TCA genes show are upregulated in Selected populations. Distribution of differences between Selected and Control flies raised on poor diet in expression of 33 genes for enzymes catalyzing glycolysis and the TCA cycle (blue), compared to 1000 random samples of other genes with the same distribution of overall mean expression levels (yellow).

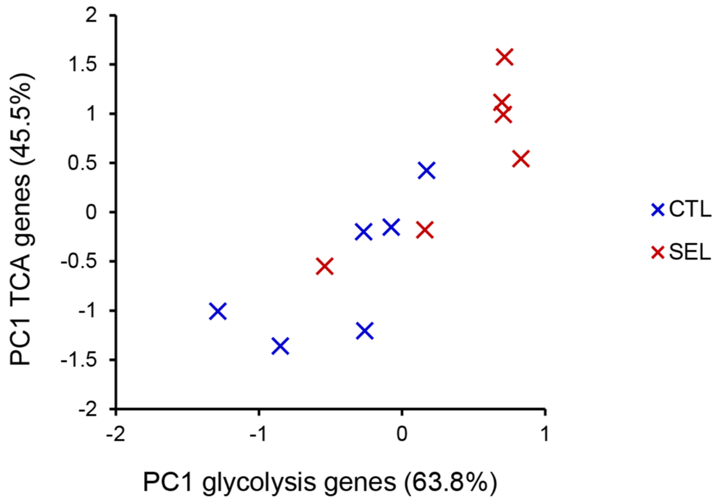


**Supplementary Figure S2**. Expression of glycolysis and TCA genes is correlated across populations. PC1 scores for the expression levels of 12 glycolysis genes and 21 genes coding for TCA enzymes for Selected and Control flies raised on poor diet, $r = 0.88$, $P = 0.0002$. Both scores are significantly different between the Selected and Control populations (glycolysis: $t_{10} = 2.8$, $P = 0.020$; TCA: $t_{10} = 2.6$, $P = 0.024$).

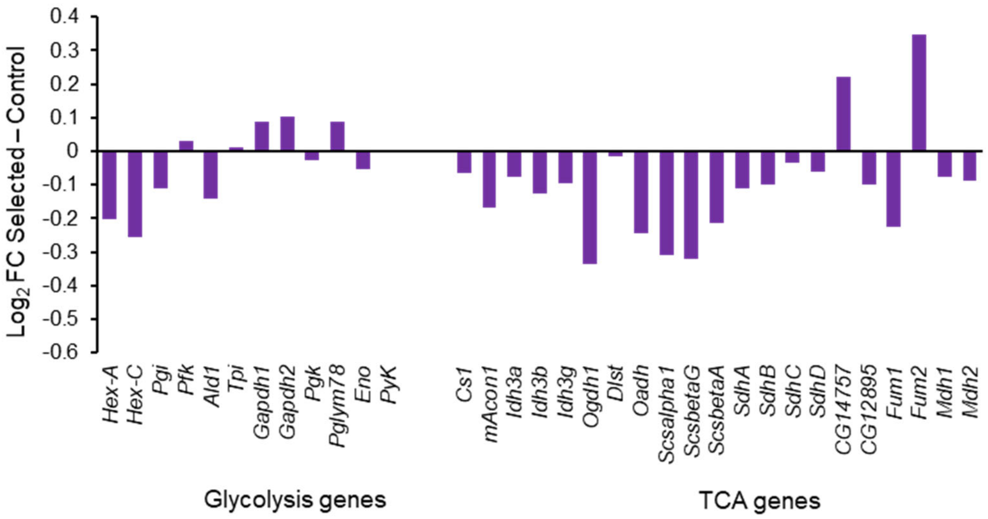


**Supplementary Figure S3.** The difference between Selected and Control populations in expression of glycolysis and TCA cycle genes in early third instar larvae raised on poor diet. Data from (Erkosar et al., 2017).

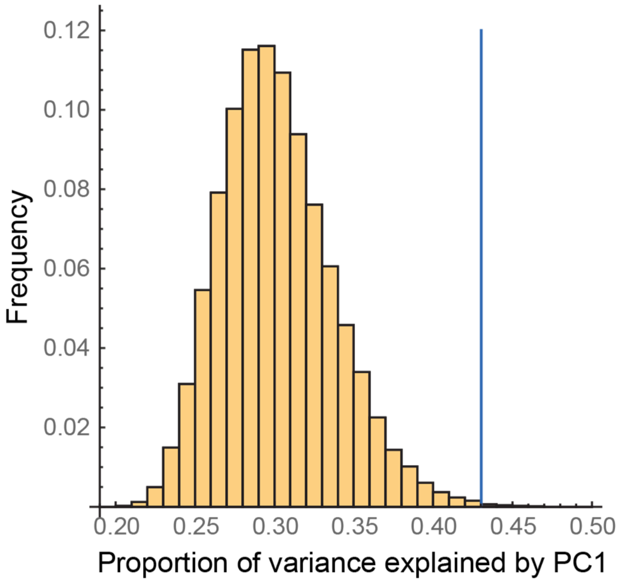


**Supplementary Figure S4.** The proportion of variance explained by the first principal component of the PCA on population residuals of nine traits from Figure 5 (blue line) compared to the distribution of the proportion of variance explained by PC1 on 100,000 randomized data sets (histogram, median = 0.30). The randomized data sets were generated by random permutations of replicate populations within evolutionary regimes separately for each trait, preserving their attribution to the Selected versus Control regime. Only a fraction of 0.0015 of randomized data sets produced a PC1 that accounted for more variance than the PC1 on the actual data.

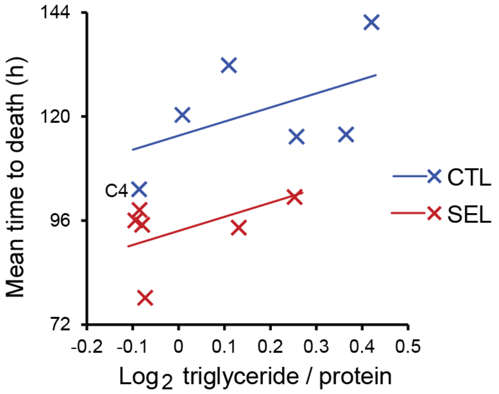


**Supplementary Figure S5.** The relationship between the mean time to death for each population and its mean triglyceride content. The lines are predictions from ANCOVA: however, the slope is not significantly different from zero ($F_{1,9}$ = 3.2, $P$ = 0.11). Control population C4 (indicated) clustered with Selected populations for both traits.

### Supplementary Tables

**Supplementary Table S1**. Primers used in the qPCR to quantify the ratio of mitochondrial to nuclear genomes.

| Gene | Forward primer | Reverse primer | Efficiency |
|---|---|---|---|
| *Cox1* | GATTAGCTACTTTACATGGAACTC | CTGCTATAATAGCAAATACAGCTC | 2.14 |
| *ATP6* | TCCTCAAGGAACACCCGCTA | ACAGCTAATGTTCCAGGTCGAA | 1.99 |
| *Actin42a_PCR* | CAGATGTGGATCTCGAAGCA | TTCTGAAGGAGCGGAAGTGT | 2.09 |
| *Rpl32_PCR* | GCCGCTTCAAGGGACAGTATCTG | AAACGCGGTTCTGCATGA | 2.02 |

**Supplementary Table S2**. Results of type 3 *F*-tests of fixed factors in the GMM of (A) protein, (B) glycogen and (C) triglyceride content (all log-transformed), and (D) estimates of variance components for the random effects in those models.

(A) Protein content per larva

| Effect | *df* | *F* | *P* |
|---|---|---|---|
| Diet (standard vs. poor) | 1,18.2 | 165.3 | **< 0.0001** |
| Stage (day 1 vs. day 4) | 1,2.4 | 0.0 | 0.95 |
| Regime (Control vs. Selected) | 1,20.5 | 5.6 | **0.0275** |
| Diet×Regime | 1,18.2 | 0.3 | 0.62 |
| Stage×Regime | 1,10.4 | 2.5 | 0.14 |
| Stage×Diet | 1,10.1 | 4.2 | 0.066 |
| Stage×Diet×Regime | 1,10.1 | 3.7 | 0.082 |

(B) Glycogen content relative to protein, full model

| Effect | *df* | *F* | *P* |
|---|---|---|---|
| Diet (standard vs. poor) | 1,10.1 | 17.8 | **0.0017** |
| Stage (day 1 vs. day 4) | 1,5.1 | 273.1 | **0.0001** |
| Regime (Control vs. Selected) | 1,10.0 | 0.0 | 0.91 |
| Diet×Regime | 1,10.1 | 0.8 | 0.38 |
| Stage×Regime | 1,10.1 | 1.3 | 0.28 |
| Stage×Diet | 1,10.1 | 0.1 | 0.81 |
| Stage×Diet×Regime | 1,10.1 | 1.2 | 0.30 |

(C) Triglyceride content relative to protein, full model

| Effect | *df* | *F* | *P* |
|---|---|---|---|
| Diet (standard vs. poor) | 1,10.8 | 63.8 | **<0.0001** |
| Stage (day 1 vs. day 4) | 1,9.4 | 0.8 | 0.39 |
| Regime (Control vs. Selected) | 1,17.3 | 15.4 | **0.0011** |
| Diet×Regime | 1,10.8 | 3.1 | 0.11 |
| Stage×Regime | 1,17.5 | 0.8 | 0.38 |
| Stage×Diet | 1,10.3 | 15.8 | **0.0025** |
| Stage×Diet×Regime | 1,10.3 | 12.3 | **0.0055** |

(D) Variance component estimates for random effects

| | Protein | | Glycogen | | Triglycerides | |
|---|---|---|---|---|---|---|
| Random effect | Estimate | SE | Estimate | SE | Estimate | SE |
| Population | 0.0000 | - | 0.0121 | 0.0092 | 0.0000 | - |
| Stage×Population | 0.0040 | 0.0075 | 0.0058 | 0.0061 | 0.0238 | 0.0125 |
| Diet×Population | 0.0472 | 0.0190 | 0.0016 | 0.0045 | 0.0003 | 0.0083 |
| Stage×Diet×Population | 0.0133 | 0.0089 | 0.0046 | 0.0059 | 0.0143 | 0.0117 |
| Batch | 0.0194 | 0.0199 | 0.0010 | 0.0017 | 0.0016 | 0.0032 |
| Residual | 0.0131 | 0.0027 | 0.0158 | 0.0033 | 0.0199 | 0.0046 |

**Supplementary Table S3.** GMM of the triglyceride content separately by stage (day 1 vs day 4); adjusted *P* refers to Tukey-adjusted *P*-values.

(A) Type 3 tests of fixed effects

| | Day 1 | | | Day 4 | | |
|---|---|---|---|---|---|---|
| Effect | df | F | P | df | F | P |
| Diet | 1,10 | 52.8 | **0.0001** | 1,10 | 16.6 | **0.0022** |
| Regime | 1,10 | 10.0 | **0.01** | 1,9.17 | 6.3 | **0.0333** |
| Diet*Regime | 1,10 | 10.2 | **0.0097** | 1,10 | 3.6 | 0.0881 |

(B) Pairwise comparisons

| | Day 1 | | Day 4 | |
|---|---|---|---|---|
| Comparison | *P* | adjusted *P* | *P* | adjusted *P* |
| CTL poor vs. SEL poor | **0.0004** | **0.0062** | 0.18 | 0.52 |
| CTL poor vs. CTL std | **0.0001** | **0.0001** | 0.15 | 0.45 |
| CTL poor vs. SEL std | **0.0001** | **0.0003** | **0.0013** | **0.0104** |
| SEL poor vs. CTL std | 0.22 | 0.60 | 0.67 | 0.97 |
| SEL poor vs. SEL std | **0.016** | 0.066 | **0.0018** | **0.0081** |
| CTL std vs. SEL std | 0.36 | 0.79 | **0.0085** | **0.0482** |

(C) Variance component estimates for random effects

| | Day 1 | | Day 4 | |
|---|---|---|---|---|
| Random effect | Estimate | SE | Estimate | SE |
| Population | 0.025 | 0.021 | 0.020 | 0.013 |
| Diet×Population | 0.025 | 0.016 | 0.002 | 0.006 |
| Batch | 0.000 | - | 0.006 | 0.011 |
| Residual | 0.022 | 0.006 | 0.016 | 0.006 |

**Supplementary Table S4.** The results of the tests and the estimates of effect sizes for the difference between Selected and Control populations in the abundance of 113 core metabolites in poor-diet raised mature females subject to 24 h of food deprivation. Separate file.

**Supplementary Table S5**. GMM on citrate synthase activity (log-transformed).

(A) F-test of fixed effects

| Effect | df | *F* | *P* |
|---|---|---|---|
| Diet | 1,19 | 47.8 | **< 0.0001** |
| Regime | 1,19 | 24.8 | **< 0.0001** |
| Diet×Regime | 1,19 | 2.2 | 0.15 |

(B) Variance component estimates for random effects

| Random effect | Estimate | SE |
|---|---|---|
| Population | 0 | - |
| Residual | 0.040 | 0.013 |

**Supplementary Table S6**. GMM on the mtDNA copy number (the ratio of mitochondrial to nuclear genomes in fly thoraxes, log-transformed).

(A) F-test of fixed effects

| Effect | *df* | *F* | *P* |
|---|---|---|---|
| Diet | 1,10 | 8.65 | **0.0148** |
| Regime | 1,10 | 3.03 | 0.1126 |
| Diet×Regime | 1,10 | 0.12 | 0.7366 |

(B) Variance component estimates for random effects

| Random effect | Estimate | SE |
|---|---|---|
| Population | 0.0032 | 0.0028 |
| Food×Population | 0.0008 | 0.0024 |
| Block | 0.0000 | |
| Residual | 0.0128 | 0.0026 |

**Supplementary Table S7**. ANCOVA on the mitochondrial to nuclear genome ratio. The interaction (*P* = 0.51) has been removed from the model.

| Effect | *df* | *F* | *P* |
|---|---|---|---|
| Regime | 1,9 | 0.1 | 0.77 |
| PC1 of glycolysis and TCA genes | 1,9 | 6.5 | 0.031 |

**Supplementary Table S8**. GMM on total activity within 36 h and the timepoint of the last movement.

(A) Test of the fixed effect

| | Total activity in the first 36 h | | | Timepoint of the last movement | | |
|---|---|---|---|---|---|---|
| Effect | df | F | P | df | F | P |
| Regime | 1,8.88 | 5.0 | 0.052 | 1,9.75 | 14.8 | 0.0034 |

(B) Variance component estimates for random effects

| | Total activity in the first 36 h | | Timepoint of the last movement | |
|---|---|---|---|---|
| Random effect | Estimate | SE | Estimate | SE |
| Population | 0.082 | 0.047 | 39.4 | 22.1 |
| Activity monitor | 0.009 | 0.014 | 2.0 | 5.0 |
| Residual | 0.242 | 0.027 | 132.3 | 14.9 |

**Supplementary Table S9.** Variance and covariance components and correlation estimates between activity and endurance.

| Component | $Log_2$ activity | Last move | Sum | Covariance | *r* |
|---|---|---|---|---|---|
| Population within regime | 0.082 | 39.400 | 36.041 | -1.720 | -0.956 |
| Residual (individual fly) | 0.242 | 132.267 | 130.502 | -1.004 | -0.177 |
| Activity monitor + Block | 0.009 | 2.038 | 1.827 | -0.110 | -0.802 |
| Total | 0.334 | 173.706 | 168.370 | -2.835 | -0.372 |

### Supplementary Methods

*Details of fly husbandry*

Our "standard" larval diet consisted of 12.5 g dry brewer's yeast, 30 g sucrose, 60 g glucose, 50 g cornmeal, 0.5 g $CaCl_2$, 0.5 g $MgSO_4$, 10 ml 10% Nipagin, 6 ml propionic acid, 20 ml ethanol, and 15 g of agar per liter of water, the poor diet contained 1/4 of the amounts of yeast, sugars and cornmeal of the standard diet. In the course of experimental evolution and for all assays reported here flies were raised at the controlled density of about 200-250 larvae per bottle containing 40 ml of standard or poor diet, at 25°C, 60-70% ambient relative humidity and 12:12 light:dark cycle. To obtain flies used in the experiments 100-200 adults were allowed to oviposit on orange juice-agar jelly overnight. Approximately 200-250 embryos were subsequently transferred to a bottle containing 40 ml of standard or poor diet and inoculated with mixed feces from adults of all populations to provide homogenized microbiota; for detailed protocol and rationale see (Erkosar et al., 2023). On the poor diet the emergence of flies from a particular bottle can be spread over 10 days, and the Selected flies emerge 2-3 days sooner than the Controls. To assure a sufficient number of adults from all populations, multiple poor food bottles were set up in a staggered manner over several days and offset to compensate for the difference in developmental time between Selected and Control populations. Flies eclosed overnight (within about 16 h) were collected around the day of peak of emergence from at least two bottles per population and diet. Samples of freshly emerged females collected this way were stored at –80°C for analysis of protein, glycogen and triglycerides on day 1. The remaining assays were performed on mature, mated and reproducing females. To obtain them, the freshly emerged flies of both sexes were transferred to new bottles with standard diet and allowed to feed and mate for 3 days before collecting the females to be used in the experiments; we refer to those flies as 4 days old females. The exception is the assay of locomotion, which (for logistic rather than scientific reasons) was done on day 6, after the flies have been maintained on standard diet for 5 days.

*Protein, triglycerides and glycogen content*

Upon collection, the flies were anesthetized with $CO_2$ and their heads were removed with a scalpel because the red eye pigment would interfere with colorimetric assays. The samples were flash-frozen in liquid nitrogen immediately after collection and stored at -80°C until protein, triglyceride and glycogen could be measured. The following protocols were adapted from (Tennessen et al., 2014)

Fly samples were first homogenized by bead beating in 250µl of TE buffer (10 mMtris,1mM EDTA, pH 8.9). To quantify protein with the Bradford assay, two technical replicates of 2 µl (flies raised on standard diet) or 5 µl (flies raised on poor diet) of the homogenate were added to 300 µl of Coomassie Plus Reagent (Thermo Scientific 23200) and incubated for 10 minutes. Absorbance at 595 nm was measured and converted to protein concentration using a standard calibration curve obtained from 5 standards with decreasing concentrations of a reference protein (0.5, 0.25, 0.125, 0.0625 and 0.00315 g/l of BSA, Sigma P5369). Different amounts of homogenate were used because of large difference in size of flies raised on the two diets. The concentrations of proteins estimated in the samples were converted in the amount of protein per fly based on the dilutions and the number of flies per sample.

Another 80 µl of the sample homogenate was used for triglyceride quantification. It was heated at 70°C for 10 minutes to inactivate enzymes that could potentially digest triglyceride. For each sample, 2 x 20 µL of the heated homogenate were isolated in new tubes. To one tube, 20 µL of triglyceride reagent (Sigma, T2449) containing lipase was added. In this tube, triglyceride reagent digested triglycerides, releasing glycerol and fatty acids. Sample in the other tube was diluted with 20 µL of TE buffer and therefore only contained the amount of glycerol that was already initially present in the homogenized sample. Glycerol standards of known concentrations were prepared by a series of 2-fold dilutions (1,0.5, 0.25, 0.125, 0.0625 g/l equivalent triolein concentration) of Glycerol standard solution (Sigma, G7793). Samples and standards were incubated 60 minutes at 37°C for digestion to occur in tubes containing lipase. After this, samples were centrifuged for 3 minutes at 7500 rpm to remove fly body debris. 30 µL of samples and standards were loaded in wells of a 96-wells plate, and 100 µL of Free Glycerol Reagent (Sigma, F6428) were added. After 5 minutes of incubation, absorbance at 540 nm was read with a Spectrophotometer plate reader. Glycerol concentrations (corresponding to the amount of triglyceride initially present in the homogenate) were determined after subtraction of blank and glycerol background values. They were converted into triolein equivalent concentrations based on the standard curve and normalized by dividing by the protein concentration in the sample.

To quantify glycogen content, the rest of the homogenate was centrifuged for 5 minutes at 7500 rpm to remove potential fly body debris. 80µl of the supernatant heated at 70°C for 10 minutes, after which they were stored at -80°C to be treated the next day. After unfreezing, samples were diluted 1:3 with TE buffer. 2 × 40µl of sample was pipetted in new tubes; 40µl of amyloglucosidase solution (1.5µL amyloglucosidase in 1mL TE buffer ,Sigma, A1602) was added (amyloglucosidase digests glycogen to glucose). The other tube was used to determine the background glucose concentration; it was diluted with 40µl of TE instead of AS. Digestion occurred in a plate sealed with Parafilm, at 37°C for 60 minutes. 100 µL Hexokinase reagent (Sigma G3293) were then added to the samples. After 15 minutes of incubation, absorbance at 340 nm was read to quantify the amount of NADH generated in the reaction. Glycogen quantities were calculated after subtraction of the absorbance of background glucose (untreated samples), using a standard curve obtained from the absorbance of known glycogen (Sigma, G0885) concentrations (1,0.5, 0.25, 0.125, 0.0625 g/l respectively). Glycogen concentrations were normalized to sample protein content. Because glycogen was quantified in centrifugated samples, we additionally repeated protein quantification in the samples after centrifugation with the same protocol as described above. The quantities of glycogen normalized to protein content before and after centrifugation were nearly perfectly correlated ($r$ = 0.996) and led to identical conclusions. We thus only report glycogen results normalized to protein content before centrifugation.

*Citrate synthase*

Fly samples were manually crushed with a pestle in 1 ml of isolation buffer (IB, 6.846g sucrose, 0.121g Tris and 1ml EGTA 0.1M in 100ml $H_2O$ pH7.4, two tablets of protease inhibitor). Protein content of each sample was measured from this homogenate following the same protocol as described above. Mitochondrial fraction was then isolated from the homogenate. First, samples were centrifued at 850 RCF for 5 minutes at 4°C, then supernatant was transferred to a new tube and centrifuged again at 1’000 RCF for 5 minutes at 4°C. Supernatant was transferred a second time and centrifuged at 13’000 RCF for 5 minutes at 4°C. The supernatant was then removed, and pellet was washed with 1ml IB. The resuspended extract was spun at 13’000 RCF for 5 minutes at 4°C, and supernatant was removed. Samples were resuspended in 30µl of PBS and stored at −80°C until citrate synthase activity measurement. Samples were diluted 2X in PBS before measurement occurred. Citrate synthase activity was measured from mitochondrial fractions using Citrate Synthase Activity Assay Kit (Sigma, #MAK193) following manufacturer’s instructions. Briefly, CS activity was determined using a coupled enzyme reaction resulting in a colorimetric product characterized by absorbance of 412 nm. Change in absorbance is proportional to the enzymatic activity. Absorbance was measured for 10 hours every 10 min at 25°c and 412 nm. To calculate the rate of reaction, we used absorbance data from 60 to 240 min (later the decline in the rate became apparent); we estimated the reaction rate during this period as the slope of regression fitted to the absorbance over time.

References

Erkosar B, Dupuis C, Cavigliasso F, Savary L, Kremmer L, Gallart-Ayala H, Ivanisevic J, Kawecki TJ, 2023. Evolution under juvenile malnutrition impacts adult metabolism and impairs adult fitness in Drosophila. ELife 12:e92465. doi: 10.7554/eLife.92465.

Erkosar B, Kolly S, van der Meer JR, Kawecki TJ, 2017. Adaptation to chronic nutritional stress leads to reduced dependence on microbiota in *Drosophila*. mBio 8:e01496-01417. doi: 10.1128/mBio.01496-17.

Tennessen JM, Barry WE, Cox J, Thummel CS, 2014. Methods for studying metabolism in *Drosophila*. Methods 68:105-115. doi: https://doi.org/10.1016/j.ymeth.2014.02.034.
