## Supplementary material for "Shared genetic architecture links energy metabolism, behavior and starvation resistance along a power-endurance axis": Original Data Files: Readme.txt

Data from Erkosar et al
"Shared genetic architecture links energy metabolism, behavior and starvation resistance along a power-endurance axis"

Data files:
Data_Protein_TAG_Glycogen.xlsx : Data on protein, triglyceride and glycogen content
Data_Starvation_Resistance.xlsx : Data on time to death under starvation
Data_Citrate_Synthase.xlsx : Reaction rates, normalized reaction rates and protein content of the sample from the analysis of enzymatic activity of citrate synthase
Data_Mitochondrial_DNA_Copy_Number.xlsx : qPCR data on genomic copies of two mitochondrial and two nuclear genes
Data_Locomotor_Activity : Data on locomotor activity of starved flies
